# Specific amygdala and hippocampal subfield volumes in social anxiety disorder and their relation to clinical characteristics – an international mega-analysis

**DOI:** 10.1101/2024.01.29.576056

**Authors:** Ziphozihle Ntwatwa, Jule M. Spreckelmeyer, Janna Marie Bas-Hoogendam, Jack van Honk, Mary M. Mufford, Carl-Johan Boraxbekk, Jean-Paul Fouche, Andreas Frick, Tomas Furmark, Heide Klumpp, Christine Lochner, K Luan Phan, Kristoffer N.T. Månsson, J. Nienke Pannekoek, Jutta Peterburs, Karin Roelofs, Annerine Roos, Thomas Straube, Henk van Steenbergen, Marie-José Van Tol, Dick J. Veltman, Nic J.A. van der Wee, Dan J. Stein, Jonathan C. Ipser, Nynke A. Groenewold

## Abstract

Social anxiety disorder (SAD) has been associated with alterations in amygdala and hippocampal volume but there is mixed evidence for the direction of volumetric alterations. Additionally, little is known about the involvement of the distinct subfields in the pathophysiology of SAD. Volumetric data from a large multi-centre sample of 107 adult individuals with SAD and 140 healthy controls (HCs) was segmented using FreeSurfer to produce 9 amygdala and 12 hippocampal subfield volumes. Volumes were compared between groups using linear mixed-effects models adjusted for age, age-squared, sex, site and whole amygdala and hippocampal volumes. Subgroup analyses examined subfield volumes in relation to comorbid anxiety disorder, and comorbid major depressive disorder (MDD), psychotropic medication status, and symptom severity. In the full sample, SAD was associated with smaller amygdala volumes in the basal (*d=-*0.32, p*_FDR_*=0.022), accessory basal (*d=-* 0.42, p*_FDR_*=0.005) and corticoamygdaloid transition area (*d=-*0.37, p*_FDR_*=0.014), and larger hippocampal volume in the CA3 (*d=*0.34, p*_FDR_*=0.024), CA4 (*d=*0.44, p*_FDR_*=0.007), dentate gyrus (*d=*0.35, p*_FDR_*=0.022) and molecular layer (*d=*0.28, p*_FDR_*= 0.033), compared to HCs. SAD without comorbid anxiety, in addition, demonstrated smaller lateral amygdala (*d=-*0.30, p*_FDR_*=0.037) and hippocampal amygdala transition area (*d=-*0.33, p*_FDR_*=0.027) relative to HCs. In SAD without comorbid MDD, only the smaller accessory basal amygdala remained significant (*d=-*0.41, p*_FDR_*=0.017). No association was found between subfield volume and medication status or symptom severity. In conclusion, we observed distinct patterns of volumetric differences across specific amygdala and hippocampal subfields, regions that are associated with sensory information processing, threat evaluation and fear generalization. These findings suggest a possible disruption in information flow between the amygdala and hippocampal formation for fear processing in SAD.

## 1. Introduction

Social anxiety disorder (SAD), previously referred to as social phobia, is a debilitating anxiety disorder characterized by excessive fear and avoidance of social situations. Typically, individuals with SAD exhibit traits of social fearfulness or experience heightened anxiety towards and during social and performance-based tasks ^1^. The social fears may lead to avoidance behaviours in daily and professional life (Stein and Stein, 2008), which in its more severe forms increases the risk of school drop-out, work absenteeism, unemployment ^3^, and functional disability ^4^. SAD has a 12-month and lifetime prevalence of 7.1% (SE=0.3) and ± 12.1% (SE=0.4) respectively (Stein and Stein, 2008). Clinical studies suggest that SAD is often comorbid with major depressive disorder (MDD) and other anxiety-related disorders, including panic disorder and other phobias, with MDD most likely to develop within the first two years after the onset of SAD ^5^.

Neuroimaging studies suggest that SAD is associated with dysregulation in the brain’s ‘fear circuit’ consisting of the amygdala, hippocampus, insula, anterior cingulate, and prefrontal cortex (PFC). While there is evidence of abnormalities in the volume of the amygdala and hippocampus, findings in often small samples have proved inconsistent as both smaller ^6,7^, larger ^8^ and no differences in volumes ^9^ have been reported in individuals with SAD compared to HCs. Large-scale studies, however, using pooled structural MRI data from various international research centres, suggest that there is no difference in the volume of amygdala and hippocampus between individuals with SAD and HCs (Bas-Hoogendam et al., 2017; Wang et al., 2021).

One possible explanation for the inconsistent findings is the influence of clinical characteristics on volumetric differences observed in subcortical regions in SAD ^12^. A recent mega-analytic study using 37 samples consisting of 1115 individuals with SAD and 2775 HCs was conducted by ENIGMA-Anxiety. This study found smaller amygdala volumes in SAD individuals with comorbid anxiety disorder and comorbid MDD ^13^. Additionally, given that the amygdala and hippocampus consist of structurally heterogenous units of specialized subfields that can be identified by their cytoarchitecture, histochemistry and connectivity profile ^14^, it is likely that investigating the volume of the amygdala and hippocampal as whole structures might not reveal subfield specific differences that may occur in SAD.

There are multiple lines of evidence that suggest the distinct roles of the amygdala and hippocampal subfields in fear, threat, and anxiety. In the human amygdala, the centromedial (CM) amygdala is suggested to be reactive to aversive outcomes but not to predictive aversive cues ^15^. On the other hand, the basolateral (BLA) amygdala is associated with threat evaluation ^16,17^ and responsive to aversive cues in high trait-anxiety ^15^. Rodent studies suggest that anxiety-like behaviours in social interactions are mediated by the activation of specific subfields, for example, activation of the BLA to central amygdala projections temporarily reduced anxiety behaviours, whilst the inhibition of this pathway had the opposing effect (Tye et al., 2011). Similarly, another rodent study found that activation of the BLA to ventral hippocampal projections reduced social behaviours, whereas inhibition of this pathway increased social behaviours ^18,19^.

Given the roles of the amygdala and hippocampus in fear and anxiety ^20,21^, and developments in amygdala and hippocampal segmentation techniques (Saygin et al., 2017), there have been increasing reports of subfield volume alterations in anxiety-related disorders ^22^. Reviews of the hippocampal subfields literature suggest that individuals with MDD have smaller volumes of the CA3/4 and larger volume of the hippocampus–amygdala transition area (HATA) compared to HCs ^23^, while individuals with PTSD have smaller volumes of the CA1/3 and dentate gyrus (DG) compared to HCs (Ben-Zion et al., 2023). Moreover, Koch and colleagues (2021) found that a smaller dentate gyrus volume prospectively predicted PTSD vulnerability. With respect to the amygdala, Morey et al. (2020) found smaller lateral and paralaminar subfields, but larger central, medical and cortical amygdala subfields in PTSD compared (n=149) to HCs. Zhang et al. (2021) found larger medial amygdala subfields, whereas all the other of the subfields were found to be smaller in PTSD (n=69) compared to HCs. Taken together, these findings suggest that subfields volumes may be differentially altered depending on the disorder.

Through collaborative international efforts we leveraged multi-site data to examine amygdala and hippocampal subfield volumes in a unique large sample of 107 adult individuals with SAD and 140 HCs. We used an *in vivo* automated segmentation algorithm on T1-weighted MRI scans, allowing for the analysis of 9 amygdala and 12 hippocampal subfields. In line with the evidence of the influence of clinical characteristics on subcortical volumes in SAD, we performed subgroup analyses that were restricted to SAD without lifetime 1) comorbid anxiety disorder, 2) comorbid MDD, and 3) psychotropic medication use. Lastly, in a post-hoc analysis, we explored whether volumetric differences in individuals with SAD are greater with higher symptom severity.

## 2. Methods

### 2.1. Participants and MRI acquisition

Sociodemographic and neuroimaging data were obtained from a previous voxel-based morphometry multi-centre mega-analysis, which was part of the European and South African Research Network in Anxiety Disorders (EUSARNAD; Baldwin & Stein, 2012; Bas-Hoogendam et al., 2017). Data had been aggregated from multiple completed studies, conducted in five different countries (Germany, Sweden, United States of America, The Netherlands, and South Africa). The sample description has been detailed in a previous publication (Bas-Hoogendam et al., 2017). Briefly, ethical approval was obtained per site from local ethical review boards and written informed consent was obtained from each participant. Participants were sourced through clinics and public announcements. All participants were assessed using either the structured clinical interview (SCID) ^26^, mini-international neuropsychiatric interview (MINI) ^27^ or composite international diagnostic interview (CIDI) ^28^. The diagnosis group were required to have SAD as the primary diagnosis, and HCs were participants without lifetime psychiatric disorders. The exclusion criteria were: age younger than 18 or older than 65 years, and presence of general MRI contraindications (ferromagnetic implants, claustrophobia, pregnancy). The data that was collected also included information on clinical characteristics such as psychiatric comorbidity (comorbid anxiety disorder and comorbid MDD), psychotropic medication use, age of onset of SAD, and SAD illness severity (as measured by the Liebowitz Social Anxiety Scale (LSAS) ^29^. Structural T1-weighted 3T MRI scans were collected from all participants (details on the scan parameters can be found in the Supplementary Materials).

### 2.2. MRI image analysis and segmentation

Image analysis was performed on the University of Cape Town’s High-performance computing (HPC) cluster, Cape Town, South Africa. First, we applied the standard Freesurfer (FS) v5.3 analysis pipeline using *recon-all* to initiate all cortical reconstruction processes (http://surfer.nmr.mgh.harvard.edu/)*. Recon-all* initiates bias-field correction to the T1-weighted images, as well as registration to Talairach space, intensity normalisation, and skull stripping ^30^.

Next, subfield segmentation was performed using the *segmentHA_T1.sh* script bundled with FS v6.0. This script simultaneously segments the amygdala and hippocampus (AH), thereby preventing structural overlap. The probability atlas applied by the script is based on the transformation of *ex vivo* manual segmentation to an automated algorithm that segments *in vivo* MRI data in target space. The atlas was built using Bayesian inference based on a tetrahedral mesh spanning the amygdala and neighbouring structures ^31^. Subfields are illustrated in Figure 1. The following amygdala subfields were extracted: lateral (LA), basal (BA), accessory basal (AB), central (Ce), medial (Me), cortical (Co), paralaminar nucleus (PL) and 2 transition areas (anterior amygdaloid area (AAA) and cortico-amygdaloid transition (CAT)). These amygdala subfields can be grouped into amygdala’s three main structures; 1) the BLA complex containing the LA, BA, AB, and PL, 2) the centro-medial containing the Ce and Me nucleus, and 3) the superficial area containing the AAA, Co, and CAT ^32^. The hippocampus was segmented into the following subfields: parasubiculum, presubiculum, subiculum, three cornu ammonis (CA) sectors (CA1, CA2-3, CA4), dentate gyrus (DG), molecular layer (ML), hippocampus–amygdala transition area (HATA), fimbria, hippocampal tail, and hippocampal fissure ^33^. In addition, we also extracted whole volumes of the AH as well as total intracranial volume (ICV).

**Figure 1:**
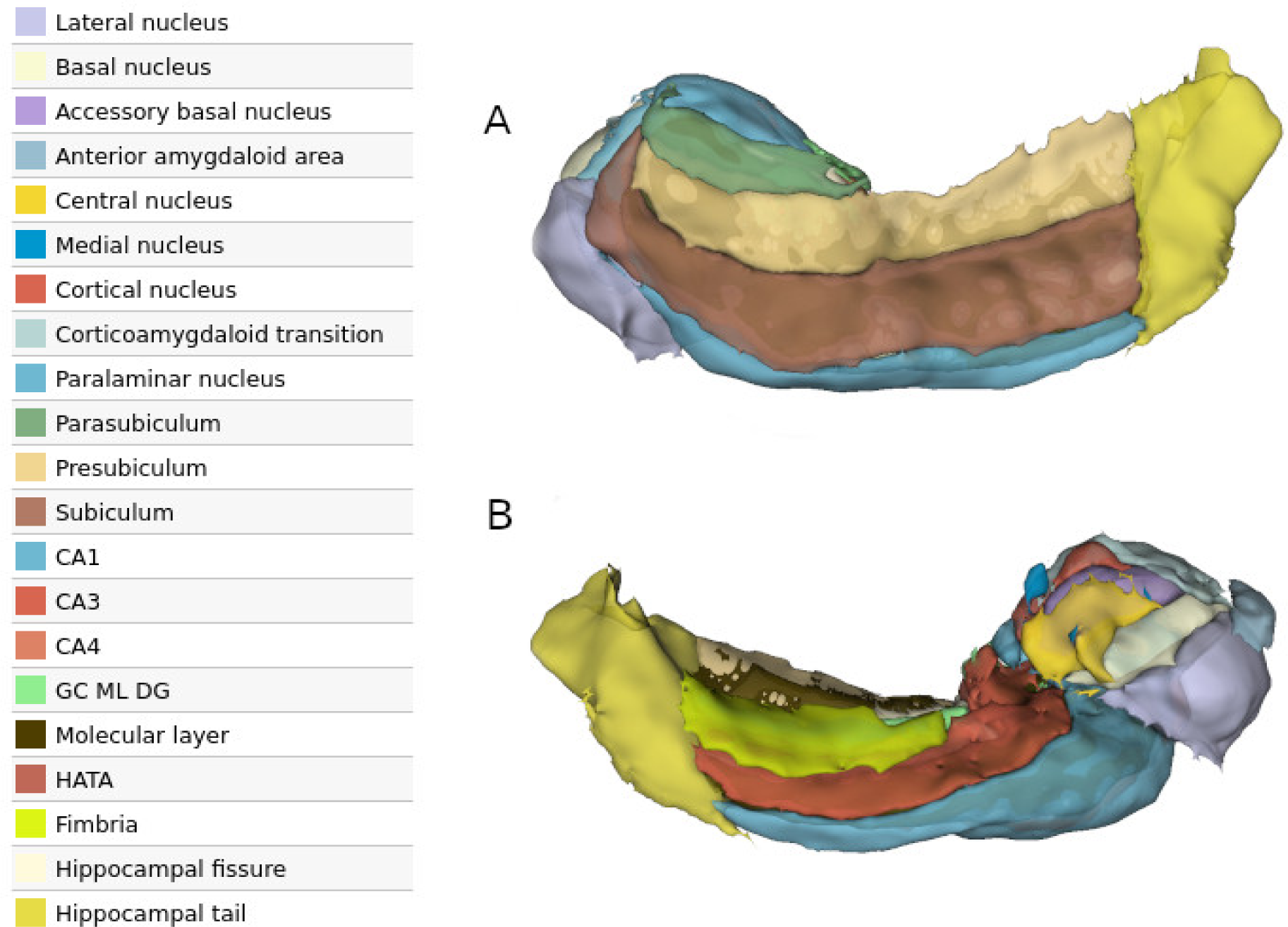
Visualisation of amygdala hippocampal Freesurfer subfield segmentation from right hemisphere of single representative (healthy control). The image was generated using 3DSlicer (https://www.slicer.org/). A: Lateral view, B: Medial view. Abbreviations: cornu ammonis (CA) sectors, CA1, CA2-3, CA4, granule cell layer of dentate gyrus (DG), molecular layer (ML), hippocampus–amygdala transition area (HATA), corticoamygdaloid transition area (CAT), anterior amygdaloid area (AAA).

### 2.3. Quality control

We used a combination of visual inspection and quantitative measures to identify inaccurate subfield segmentation. To this end, we used an adaptation of the Enhancing Neuroimaging Genetic through Meta-Analysis (ENIGMA) ^34^. Quality Control protocols for subcortical brain regions and hippocampal subfields (see https://enigma.ini.usc.edu/protocols/imaging-protocols/). In brief, each T1-weighted scan was examined by two independent raters, ZN and JMS, who were blind to the diagnostic status of the participant. Scans were examined for poor quality including motion and other scan artefacts, insufficient contrast, or presence of anatomical deviations, and segmentations were examined for incorrect assignment of the subfields ^35^. The Freeview utility included with FreeSurfer was used for 3D inspection to confirm that poor quality scan data did not meet quality standards for inclusion. Any subjects for whom the same subfield was identified as problematic through visual QC and outlier detection procedures were excluded from subsequent analyses.

### 2.4. Statistical analysis

#### 2.4.1. Covariate selection

Covariates were selected based on their established association with amygdala and hippocampal volume. We corrected for age, age-squared, sex, and scanner site (Barnes et al., 2010; Chen et al., 2018; Nordenskjöld et al., 2013; Sargolzaei et al., 2015; and consistent with Groenewold et al., 2023). We also corrected for total subfield volume using the combined AH rather than ICV, as recommended by the developers of FreeSurfer (https://surfer.nmr.mgh.harvard.edu/fswiki/HippocampalSubfields). SAD can contribute to lower educational achievement (Stein and Stein 2008), and furthermore, we did not find a significant difference in education between individuals with SAD and HCs, and we thus did not use education as a covariate in our analysis.

#### 2.4.2. General Linear Models

All statistical analyses were conducted in R (https://www.r-project.org/). As we did not have an a-priori hypothesis regarding effects of laterality on subfield volumes, we combined the volumes of the left and right hemisphere to produce a single bilateral value per subfield per participant ^40^. However, we performed a post-hoc analysis to assess possible hemispheric differences in AH subfield volumes ^41^ between individuals with SAD and HCs (Supplementary Materials). In our main analysis, the bilateral AH subfield volumes were used as dependent variables in separate models, with diagnostic group (HC, SAD) as the main independent variable and scanner site included as random intercept. In total, tests were performed for 21 separate subfields. Analyses were corrected for multiple comparisons across all 21 subfields using the false discovery rate (FDR)((Benjamini & Hochberg, 1995)). We used the R package *lme4* with restricted maximum likelihood (ReML) ^43^ and outputted mixed effect (*d*) sizes, as calculated using the t values from linear mixed effects models (equation 22, Nakagawa & Cuthill, 2007). We performed separate subgroup analyses where we investigated the association between AH subfield volumes and clinical characteristics. Here, we selected participants with SAD, leaving out those with lifetime 1) comorbid anxiety disorder (SAD=86), 2) comorbid MDD (SAD=83), and 3) medication use (SAD=89), compared to HC (n=140). Participants with SAD that had unknown comorbidities were retained in this analysis. Due to small sample sizes and collinearity between comorbidities and scan sites we did not perform subgroup analyses for individuals with SAD with 1) comorbid anxiety disorder (SAD=21), 2) comorbid MDD (SAD=24), and 3) medication use (SAD=18) (for sample description see supplementary table S6). Last, we assessed the association between AH subfield volumes and SAD symptom severity (as measured by the LSAS).

## 3. Results

### 3.1. Sample characteristics

In the full sample (SAD: n=107; HC: n=140), we found no significant difference in age (SAD: 32.21 years; HC: 34.72 years; t=1.86, p-value=0.06), sex (SAD: 42% male; HC: 62%; t=0.79, p-value=0.42) and mean years of education (SAD: 14.64 years; HC: 15.12 years; t=0.95, p-value=0.34), between individuals with SAD and HCs (see Table 1). Additionally, we found no significant difference in the whole hippocampal volume between SAD and HCs (SAD: 3504.92; HC: 3472.43; t=-0.74, p-value=0.45). However, the whole amygdala volume was found to be significantly smaller in individuals with SAD compared to HCs (SAD: 1814.73; HC: 1873.26; t=2.23, p-value=0.02).

**Table 1:** SAD and HC sample composition as included in the main analysis.

|  |  | Initial number of scans <sup>a</sup> |  |  | Excluded after visual QC |  |  | Included number of scans |  |  |
| --- | --- | --- | --- | --- | --- | --- | --- | --- | --- | --- |
| Country | Research site | SAD | HC | Total | Segmentation fail | Visual QC + outlier | Missing data | SAD | HC | Total |
| The Netherlands | VU Medical Centre Amsterdam<br>NESDA Study | 6 | 27 | 33 | 0 | 3 | 0 | 6 | 24 | 30 |
|  | University of Groningen<br>NESDA Study | 8 | 11 | 19 | 0 | 0 | 0 | 8 | 11 | 19 |
|  | Leiden University Medical Centre<br>NESDA Study | 8 | 26 | 34 | 0 | 4 | 2 | 7 | 23 | 30 |
|  | Leiden University Medical Centre-Social Anxiety Study | 20 | 20 | 40 | 1 | 2 | 0 | 20 | 17 | 37 |
| South Africa | University of Cape Town;<br>Stellenbosch University | 12 | 11 | 23 | 0 | 5 | 4 | 9 | 9 | 18 |
| Sweden | Umeå University | 26 | 23 | 49 | 0 | 0 | 0 | 26 | 23 | 49 |
| Germany | University of Muenster | 19 | 22 | 41 | 0 | 0 | 0 | 19 | 22 | 41 |
| **United States of America** | University of Illinois | 12 | 11 | 23 | 0 | 0 | 0 | 12 | 11 | 23 |
| **Total** |  |  |  |  |  |  | 6 | 107 | 140 |  |

<sup>a</sup>Reported in previous publication <sup>10</sup>. <sup>b</sup> Participants excluded based on visual screening for partial or atypical segmentation using an adaptation of the ENIGMA Consortium Quality Control protocol for subcortical and hippocampal subfield (see https://enigma.ini.usc.edu/ongoing/enigma-hippocampal-subfields/).

**Table 2A:** Demographic characteristics of SAD and HC participants.

|  | SAD (n=107) |  | HCs (n=140) |  | Statistical analysis |  |  |
| --- | --- | --- | --- | --- | --- | --- | --- |
| Demographics | mean | SD | mean | SD | t | df | p |
| Age (years) | 32.2 | 10.43 | 34.3 | 10.5 | 1.86 | 229.18 | 0.06 |
| Education level (years) | 14.7 | 3.0 | 15.2 | 3.0 | 0.95 | 133.42 | 0.34 |
|  | **N** | **%** | **N** | **%** |  |  |  |
| Male | 42 | 39.25 | 62 | 44.3 | -0.79 | 230.05 | 0.43 |
| Clinical characteristics | mean | SD | mean | SD |  |  |  |
| LSAS score <sup>a</sup> | 78.1 | 19.5 | -- | -- |  |  |  |
| Age at onset (years) <sup>b</sup> | 14.8 | 7.1 | -- | -- |  |  |  |
| Total amygdala volume | 1817.73 | 202.80 | 1873.26 | 205.92 | 2.23 | 229.98 | 0.02 |
| Total hippocampal volume | 3504.92 | 347.11 | 3472.43 | 331.78 | -0.74 | 222.85 | 0.45 |

<sup>a</sup> Liebowitz Social Anxiety Scale (LSAS). <sup>b</sup> measured using LSAS.

**Table 2B:** Demographic and clinical characteristics of subgroup individuals with SAD.

| | Effect size (mixed-effects $d$) | Standard Error (SE) | P value (corrected ($p_{FDR}$)) | P value (uncorrected) |
| --- | --- | --- | --- | --- |

|  | A: SAD without comorbid anxiety disorder (n=86) | B: SAD without comorbid MDD (n=83) | C: SAD without psychotropic medication (n=89) | D: HCs (n=140) |
| --- | --- | --- | --- | --- |
|  | Mean (SD) | Mean (SD) | Mean (SD) | Mean (SD) |
| **Age (years)** | 31.14 (9.81) | 31.04 (10.27) | 31.34 (10.28) | 34.72 (10.51) |
| **Education level (years)** | 14.85 (2.77) | 15.02 (2.65) | 14.71 (3.02) | 15.12(3.01) |
|  | N (%) | N (%) | N (%) | N (%) |
| **Male** | 35 (40.69) | 34 (40.96) | 34 (38.20) | 62 (44.28) |

**Note:** mean age of SAD without; (A) comorbid anxiety disorder (n=86), (B) comorbid MDD (n=83), (C) psychotropic medication (n=89), and (D) HCs (n=140).

### 3.2. Group differences

Between group comparisons were conducted on 107 individuals with SAD and 140 HCs (full aggregated sample). With regard to amygdala subfields, individuals with SAD had significantly smaller BA (*d= -*0.32, p*_FDR_*= 0.022), AB (*d= -*0.42, p*_FDR_*= 0.005) and CAT nucleus (*d= -*0.37, p*_FDR_*= 0.014) compared to HCs (after adjustment for sex, age, age-squared, and total volume of AH, in this and subsequent analyses). In contrast, the hippocampal subfields were significantly larger for the CA3 (*d=* 0.34, p*_FDR_*= 0.024), CA4 (*d=* 0.44, p*_FDR_*= 0.007), DG (*d=* 0.35, p*_FDR_*= 0.022) and ML (*d=* 0.28, p*_FDR_*= 0.033) in individuals with SAD in comparison to HCs (Figure 2A/Table 3). A trend was observed for smaller HATA between individuals with SAD and HCs; however, this did not reach the threshold for significance after correction for multiple comparison (*d*=0.27, p*_FDR_*= 0.052). In follow-up exploratory tests of laterality effects for the bilateral subfields between SAD and HCs, trends for a lateralized group difference (p*_FDR_*=0.05-0.1) were detected for some of the amygdala subfields, however none reached significance after correction for multiple comparison (TableS1 in Supplementary).

**Figure 2.**
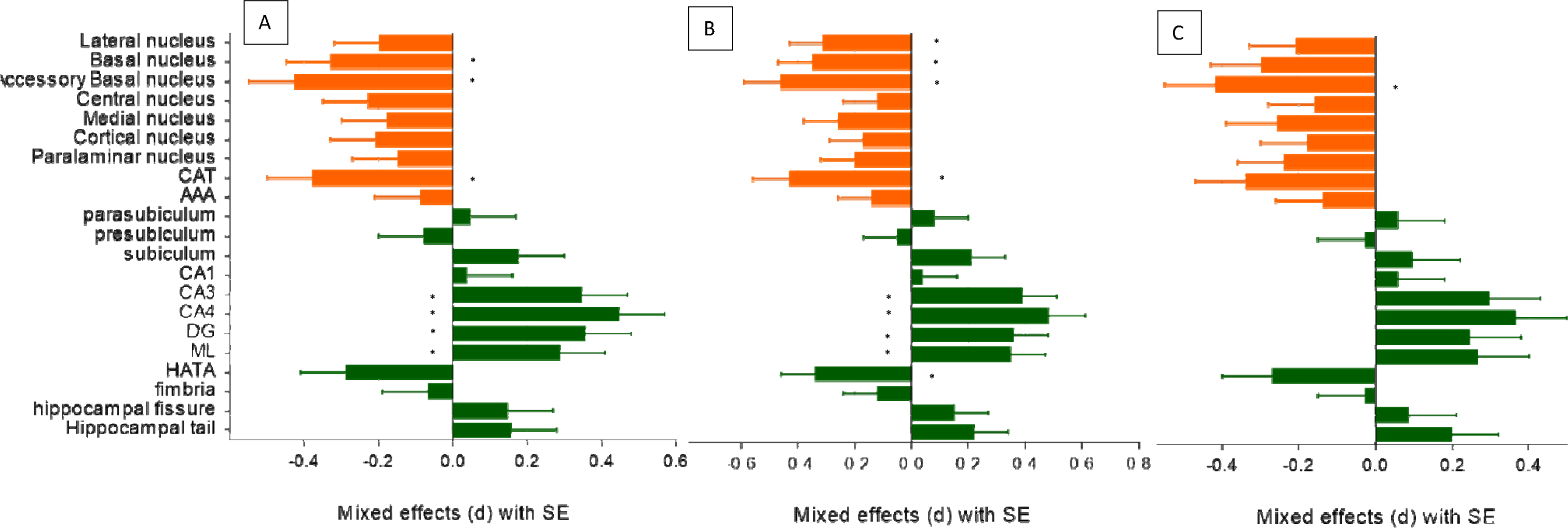
: **(A)** Mixed effect size estimates (d) for amygdala and hippocampus subfields between **individuals with SAD (n=107) and HCs (n=140)**. **(B)** Mixed effect size estimates (d) for amygdala and hippocampus subfields between **individuals with SAD without comorbid anxiety (n=86) and HCs (n=140)**. **(C)** Mixed effect size estimates (d) for amygdala and hippocampus subfields between **individuals with SAD without comorbid MDD (n=83) and HCs (n=140)**. Data presented with SE. (*) Denotes significant after FDR correction. Amygdala subfields presented in orange; hippocampal subfields presented in green. Abbreviations: cornu ammonis (CA) sectors, CA1, CA2-3, CA4, granule cell layer of dentate gyrus (DG), molecular layer (ML), hippocampus–amygdala transition area (HATA), corticoamygdaloid transition area (CAT), anterior amygdaloid area (AAA).

**Table 3:**
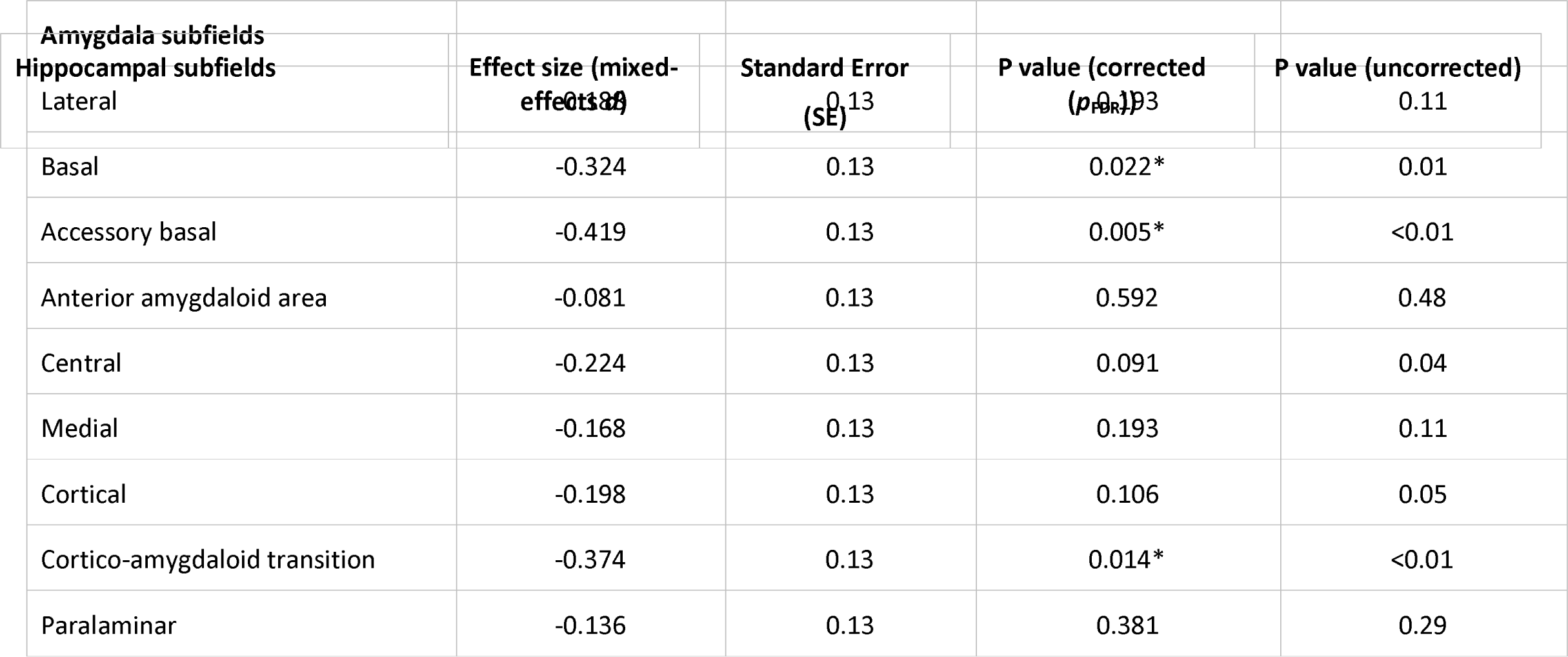
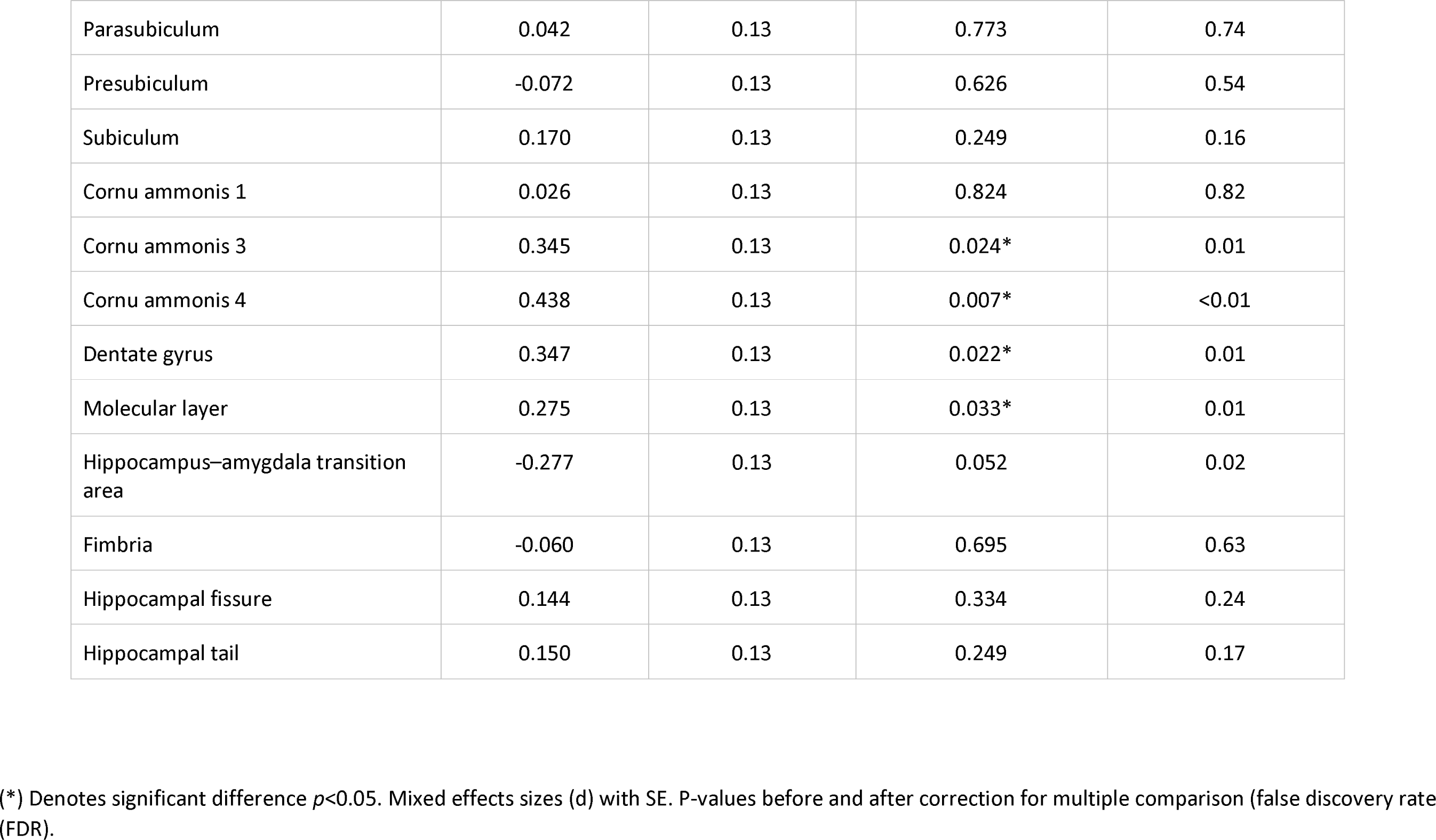
Mixed effect size estimates (d), SE, uncorrected and corrected (FDR) p-value for amygdala and hippocampal subfield volumes between individuals with SAD (n=107) and healthy controls (n=140).

### 3.3. Subgroup analyses: subfields in relation to clinical characteristics

#### 3.3.1. SAD without comorbid anxiety disorder

Individuals with SAD without comorbid anxiety disorder (n=86) were found to have significantly smaller LA and HATA subfields, in addition to those subfields that were significant in the main analysis. We observed significantly smaller LA (*d= -*0.30, p*_FDR_*= 0.037), BA (*d= -*0.34, p*_FDR_*= 0.020), AB (*d= -*0.45, p*_FDR_*= 0.005) and CAT nucleus (*d= -*0.42, p*_FDR_*= 0.007) in individuals with SAD compared to HCs (n=140). For the hippocampus, with exception of the HATA (*d= -*0.33, p*_FDR_*= 0.027), we observed significantly larger CA3 (*d=* 0.38, p*_FDR_*= 0.020), CA4 (*d=* 0.47, p*_FDR_*= 0.007), DG (*d=* 0.35, p*_FDR_*= 0.023), and ML (*d=* 0.34, p*_FDR_*= 0.018) in individuals with SAD compared to the HCs. All nuclei had generally medium effect sizes with the CA4 having the largest effect (*d=* 0.47) (Figure 2B).

#### 3.3.2. SAD without comorbid MDD

When selecting individuals with SAD without comorbid MDD (n=83), we observed that only the volume of AB nucleus of the amygdala remained significantly smaller (*d=* -0.41, p*_FDR_*= 0.017) in individuals with SAD compared to HCs (Figure 2C).

#### 3.3.3. SAD without psychotropic medication

There was no significant difference between individuals with SAD without medication use (n=89) and HCs (n=140) following adjustment for multiple comparisons (TableS4 in supplementary).

#### 3.3.4. SAD symptom severity

We did not observe any associations between symptom severity (measured with the LSAS) and amygdala nor hippocampal subfield volumes.

## 4. Discussion

The present study is the first to explore neuroanatomical differences in amygdala and hippocampal subfields related to SAD, in a large sample of adult individuals with SAD (n=107) and HC (n=140). In our primary analysis we demonstrated that individuals with SAD have significantly smaller amygdala subfields (BA, AB, and CAT) and significantly larger hippocampal subfields (CA3, CA4, DG, and ML) than HCs. In our subgroup analyses, individuals with SAD without comorbid anxiety disorder had significantly smaller LA and HATA subfields than HCs, in addition to the significant subfields in our main analysis. In individuals with SAD without comorbid MDD, only the AB amygdala remained significantly smaller compared to HCs. We did not find an association between subfield volume and psychotropic medication use or symptom severity. While previous small-scaled studies in amygdala and hippocampal volumes in SAD have yielded inconsistent results, we suggest that SAD is associated with distinct volumetric differences in specific subfields that are implicated in sensory information, threat evaluation, and pattern separation and completion. Additionally, the results suggest that subfield volume alterations are influenced by clinical characteristics, possibly contributing to the previous inconsistent findings.

### 4.1. Main analysis (SAD vs HCs): The potential involvement of individual subfields in SAD

Our main findings indicate that individuals with SAD have a smaller bilateral BLA amygdala, specifically in BA and AB subfields, compared to HCs. Post-hoc unilateral analyses showed attenuated differences, with slightly stronger effect sizes for the left compared to right amygdala. Smaller BA and AB volumes have been reported in other anxiety-related disorders with similar or smaller sample sizes as the present study (panic disorder (n = 38): Asami et al., 2018;, and PTSD: Morey et al., 2020 (n=149) ; Zhang et al., 2021 (n=69)). In rodent studies, smaller BLA volume is associated with increased fear response to auditory tones and contextual stimuli ^48^. Considering that the BA, together with the LA and AB nucleus, forms part of the primary sensory input area of the amygdala (LeDoux., 2007), and that the BA is associated with threat evaluation of sensory stimuli ^49^, it is possible that BA and AB structure could impact amygdala-dependent responses to sensory information, resulting in heightened fear and anxiety in SAD. Here, elaborate and sustained anxious appraisals as subserved by the left amygdala could be of particular interest (as discussed in Ocklenburg et al. 2022, Groenewold et al., 2023). Rodent studies suggest that the BLA dually modulates social interactions and anxiety-related behaviours through its projections to downstream targets, thereby controlling an anxiogenic phenotype ^19,50,51^. In humans, a study investigating amygdala reactivity during social-evaluative learning found hyperactivity in the basolateral amygdala in socially anxious families, suggesting a possible neurobiological endophenotype of SAD ^52^.

The BLA also projects to various structures of the neighbouring hippocampus including the trisynaptic circuitry made of the CA1, CA3 and DG ^53^. BLA-hippocampal projections have been shown to influence social interactions in rodent studies ^19^. In our study we observed significantly larger hippocampal subfield volume in the CA3, CA4, DG and ML, in SAD compared to HCs. These findings are contrary to what has been reported in other anxiety disorders, as smaller CA3/4 and DG were found in PTSD compared to HCs (Hayes et al., 2017; Postel et al., 2021; Wang et al., 2010). A study comparing hippocampal subfields in PTSD to SAD, in the context of childhood trauma, found that only the parasubiculum and HATA were smaller in PTSD ^56^. However, this study used smaller sample sizes (n<30) than the present study, and individuals with SAD were not compared to HCs. While our results indicate a trend towards smaller HATA in SAD compared to HCs, this trend did not reach significance in the main analysis.

A possible explanation for hippocampal volume differences (CA3, CA4, DG, and ML) observed in our study may be neuroplasticity and neurogenesis, which are processes that are supported by the CA4 and DG ^53,57^. Studies suggest that adult hippocampal neurogenesis modulates DG-associated pattern separation, which in turn influences the overgeneralization of fear ^58,59^. Deficits in pattern separation and irregular fear generalization may lead to the inability to differentiate between safe and unsafe stimuli, thereby producing exaggerated fear and anxiety responses to innocuous social stimuli ^58^. However, it is unclear whether hippocampal subfield volume differences reflect an adaptive response to heightened anxiety through neuroplasticity, or if it is a pre-existing risk factor for SAD, or both. Further investigation ideally using prospective longitudinal designs is required to fully elucidate this relationship.

### 4.2. Subgroup analyses (clinical characteristics)

#### 4.2.1. SAD without comorbid anxiety disorder

In subsequent analyses on subgroups of individuals with SAD without comorbid anxiety disorder (n=86), we observe that in addition to the subfields observed in our main analysis (BA, AB, CAT, CA3, CA4 DG and ML), the LA and HATA were significantly smaller in individuals with SAD without comorbid anxiety disorder, compared to HCs. This suggests that comorbid anxiety disorders may have masked volumetric differences resulting in non-significant findings for the LA (main analysis *d*=-0.18; without comorbid anxiety disorder *d*=-0.30) and HATA (marginally significant) in the main analysis. Our findings are contrary to Groenewold et al. (2023) who found smaller amygdala volumes in SAD individuals with comorbid anxiety disorder, but no differences in individuals without comorbid anxiety disorder ^13^. Methodological differences may partly account for the discrepancy, as the present study investigated subfield volumes relative to the whole amygdala volume whereas Groenewold et al. (2023) focused on the whole amygdala volume.

Of note, the subfields where the differences remained significant (BA, AB, CAT, CA3, CA4 DG and ML) in individuals with SAD without comorbid anxiety displayed similar effect sizes (range: 0.34-0.47) as in the main analysis (range: 0.27-0.43), which suggests that the volumes of these subfields may not be as affected by comorbid anxiety disorder. Our findings highlight the influence of clinical characteristics on subfield volumes observed in SAD. The HATA is suggested to be a connection for communication between the amygdala and hippocampus (Foo et al.,2017), and other work has indicated that the HATA may influence amygdala responses to memories stored in the hippocampus. Furthermore, the LA is thought to play a role in processing sensory information and fear conditioning. We speculate that LA and HATA dysregulation could disrupt information flow between the hippocampus and adjoining amygdala, thus altering emotional and social processing

#### 4.2.2. SAD without comorbid MDD

In individuals with SAD without comorbid MDD (n=83), only the AB subfield remained significantly smaller compared to HCs. This might partly be explained by loss of power, since the effect sizes of the relevant subfields were moderately attenuated (range: 0.20-0.41) and overlapped with those from the main analysis (range: 0.27-0.43). Of note, the applied FDR correction uses an adaptive threshold for significance that varies across sub-analyses. Studies of amygdala subfield volumes in MDD suggest that the lateral and anterior amygdaloid areas are smaller in individuals with MDD compared to HCs (individuals with SAD=147; HCs=144), whilst other studies report null findings (patient=76; HC=77). The present findings of larger hippocampal subfield volumes further contrast with recent meta-analyses which suggest that MDD is associated with smaller hippocampal CA3, CA4 and larger HATA volumes, compared to HCs. The fact that only the AB remained significant suggests that major depression comorbidity may have an influence on subfield volumes in SAD. While the sample of individuals with SAD with MDD was small (n=24), it is possible that SAD-related differences in subfield volumes are more pronounced in this particular subgroup.

#### 4.2.3. SAD without psychotropic medication use

While we did not find a significant difference between individuals with SAD without medication use (n = 89) and HCs, we observed comparable effect sizes (range 0.25-0.36), but in a smaller sample size than the main analysis. This suggests that the difference in the findings were mainly attributable to loss of effect size in the AB subfield, which no longer met the threshold for significance. A voxel-based morphometry meta-analysis on grey matter volume alterations in SAD found that the data on clinical characteristics like medication use is inconclusive as mixed effects on grey matter volume are observed in individuals with SAD (Wang et al., 2021). In our study, only a small number of individuals with SAD were taking medication at the time of scan (n= 18), thus restricting our ability to investigate this further.

#### 4.2.4. Symptom severity

Consistent with Groenewold et al. (2023) and Jayakar et al. (2020) investigating total amygdala volumes in SAD, we did not find an association between symptom severity and amygdala subfield volumes. We do note that the symptom severity data was limited and therefore we had less statistical power compared to the main analysis. Future studies with robust SAD symptom severity data, including multiple dimensions of anxious symptomatology, might be able to shed light on the possible association between severity and amygdala and hippocampal subfield volumes.

## 5. Strengths and limitations

Our study has multiple strengths but also limitations. We report on a large, aggregated sample and performed detailed subgroup analyses by investigating individuals with SAD without comorbid anxiety, without MDD, and without medication use. This allowed for observations across different clinical subgroups. However, because we used inherited data (original study Bas-Hoogendam et al., 2017) with small sample sizes for individuals with SAD with comorbid anxiety (n=21), comorbid MDD (n=24), and with medication use (n=18), we were unable to explore subfield differences in these groups. These low rates of comorbidity are the consequence of the inclusion and exclusion criteria of the original studies. Additionally, due to limited information available from the original studies, we were unable to explore other possible relevant influences of hippocampal subfield volumes, e.g., childhood adversity (as shown in other studies. Lastly, the caveat for automated segmentation is that studies show relatively low test-retest reliability for certain amygdala and hippocampal subfields, particularly the paralaminar nucleus and hippocampal fissure (compared to other subfields), both of which were included in our analysis. Further prospective longitidudinal studies are required to investigate subfield volumes in SAD over time, including in distinct clinical subgroups.

## 6. Conclusion

Our study demonstrates a distinct profile of subtle volumetric differences subfield volumes in individuals with SAD compared to HCs. Specifically, we found differences in the volumes of subfields that are associated with sensory input and fear/threat evaluation (BLA/AB) in line with previous clinical and preclinical work, as well as in pattern separation (DG) and completion (CA-3), and neurogenesis (DG and ML). Our findings further indicate that clinical characteristics are possible predictors of the volumetric characteristics of certain subfields, specifically those associated with the flow of sensory information between the amygdala and adjoining hippocampus, as noted by smaller LA and HATA in individuals with SAD without comorbid anxiety compared to HCs. The evidence of varying subfield volumes observed in SAD in the present study suggests that the heterogeneous nature of the amygdala and hippocampus may have contributed to the inconsistent volumetric findings reported for these structures in the SAD literature until now, in which the amygdala and hippocampus were considered as a whole. More research, and in particular longitudinal analysis, is required to investigate how amygdala and hippocampal subfield volumes are associated with SAD comorbid with MDD, and whether SAD-related differences in subfield volumes may be influenced by other clinical characteristics such as medication status.

## Supporting information

SAD_supplemental material

## Acknowledgments

Ziphozihle Ntwatwa was supported by the South African National Research Foundation. Janna Marie Bas-Hoogendam received grants from the Dutch Research Council (NWO; Rubicon grant 019.201SG.022), Medical Delta (Talent acceleration grant) and the Dutch Research Agenda (NWA-NeuroLabNL: Small Projects for NWA routes; NWA.1418.22.025). Henk van Steenbergen was supported by a grant from Dutch Research Council (NWO) to Bernhard Hommel. Henk van Steenbergen, J. Nienke Pannekoek and Jean-Paul Fouche were partially supported by the EU7th Framework Marie Curie Actions International Staff Exchange Scheme grant ‘European and South African Research Network in Anxiety Disorders’ (EUSARNAD). Münster (Jena) collaborators were partially supported by the Collaborative Research Center “Fear, Anxiety, and Anxiety disorders” in Münster, funded by the German Research Society (SFB/TRR-58, project C07 awarded to Thomas Straube) and by the Research Group “Person Perception” in Jena, funded by the German Research Society (grant number STR 987/6-1 to Thomas Straube). The infrastructure for the Netherlands Study of Depression and Anxiety (NESDA) was funded through the Geestkracht programme of the Netherlands Organization for Health Research and Development (ZonMw, grant number 10-000-1002) and is supported by participating universities and mental health care organizations (VU University Medical Center, GGZ inGeest, Arkin, Leiden University Medical Center, GGZ Rivierduinen, University Medical Center Groningen, Lentis, GGZ Friesland, GGZ Drenthe, IQ Healthcare, Netherlands Institute for Health Services Research (NIVEL) and Netherlands Institute of Mental Health and Addiction (Trimbos Institute)). Studies in Umea and Uppsala were supported by the Swedish Research Council and the Swedish Research Council for Health, Working Life and Welfare and the Swedish Brain Foundation. This work was made possible in part by a grant from Carnegie Corporation of New York. The funding sources had no involvement in writing this paper nor in the decision to submit this work for publication. We gratefully acknowledge the contribution of Henk Cremers in collecting data for the LUMC sample and making this available for analysis. We would also like to acknowledge the High-performance computing (HPC) cluster at the University of Cape Town, South Africa.

## Declarations

No conflicts of interest

## Contributions

Z. Ntwatwa, J. van Honk, J. Ipser, and N. Groenewold designed this study of subfield volumes in social anxiety disorder. The data was acquired by the MEGASAD collaborators and curated by J.M. Bas-Hoogendam, H. van Steenbergen and J.N. Pannekoek. The MEGASAD collaboration was funded by D. Stein and N. van der Wee. Data analysis was performed by Z. Ntwatwa, J. Ipser and N. Groenewold. Data quality control was performed by Z. Ntwatwa, J. Spreckelmeyer, and N. Groenewold. The data was interpreted by Z. Ntwatwa, J. Spreckelmeyer, M. Mufford, J. Ipser, and N. Groenewold. The draft article was written by Z. Ntwatwa, J. Spreckelmeyer, J.M. Bas-Hoogendam, D.J. Stein, J. Ipser, and N. Groenewold and all authors critically reviewed and revised this draft. All authors approved the final version of the manuscript to be published, and no other individuals have made substantial contributions that are not listed as authors. We have disclosed all conflicts of interest.

## Data availability

The sample described in this work was aggregated prior to the initiation of the ENIGMA-Anxiety Working Group, and has been contributing to ENIGMA-Anxiety. ENIGMA-Anxiety is open to sharing data from their participating samples to researchers for secondary data analysis. To request access to volumetric, clinical, and demographic data, an analysis plan can be submitted to the ENIGMA-Anxiety Working Group (http://enigma.ini.usc.edu/ongoing/enigma-anxiety/). Data access is contingent on approval by PIs from contributing samples.

## Notes

### Competing Interest Statement

The authors have declared no competing interest.

### Summary of Updates

Author list updated.

