## Supplementary material for "Specific amygdala and hippocampal subfield volumes in social anxiety disorder and their relation to clinical characteristics – an international mega-analysis": SAD_supplemental material

**Supplementary table S1a:** Mega analysis linear regression results for **bilateral** amygdala and hippocampal subfield volumes for **patients with SAD** (n=107) compared with healthy controls (HCs) (n=140), adjusting for age, sex, scan site and whole AH volumes.

|  | **Mixed effect size estimates d**  **(SAD vs HCs)** | **Standard error of *d*** | **FDR corrected**  **p value** | **Uncorrected p.value** |
| --- | --- | --- | --- | --- |
| Lateral | -0.188 | 0.128 | 0.193 | 0.110 |
| Basal | -0.324 | 0.129 | 0.022 | 0.005 |
| Accessory Basal | -0.419 | 0.129 | 0.005 | 0.000 |
| AAA | -0.081 | 0.128 | 0.592 | 0.480 |
| Central | -0.224 | 0.128 | 0.091 | 0.039 |
| Medial | -0.168 | 0.128 | 0.193 | 0.109 |
| Cortical | -0.198 | 0.128 | 0.106 | 0.050 |
| Corticoamygdaloid transition | -0.374 | 0.129 | 0.014 | 0.002 |
| Paralaminar | -0.136 | 0.128 | 0.381 | 0.291 |
| Parasubiculum | 0.042 | 0.128 | 0.773 | 0.736 |
| Presibiculum | -0.072 | 0.128 | 0.626 | 0.536 |
| Subiculum | 0.170 | 0.128 | 0.249 | 0.162 |
| CA1 | 0.026 | 0.128 | 0.824 | 0.824 |
| CA3 | 0.345 | 0.129 | 0.024 | 0.007 |
| CA4 | 0.438 | 0.129 | 0.007 | 0.001 |
| GC ML DG | 0.347 | 0.129 | 0.022 | 0.005 |
| Molecular layer | 0.275 | 0.128 | 0.033 | 0.011 |
| HATA | -0.277 | 0.128 | 0.052 | 0.020 |
| Fimbria | -0.060 | 0.128 | 0.695 | 0.628 |
| Hippocampal fissure | 0.144 | 0.128 | 0.334 | 0.239 |
| Hippocampal tail | 0.150 | 0.128 | 0.249 | 0.166 |

Abbreviations: cornu ammonis (CA) sectors, CA1, CA2-3, CA4, granule cell layer of dentate gyrus (DG), molecular layer (ML), hippocampus–amygdala transition area (HATA), corticoamygdaloid transition area (CAT), anterior amygdaloid area (AAA).

**Supplementary table S1b:** Mega analysis linear regression results for **separate** **left and right** amygdala and hippocampal subfield volumes for **patients with SAD** (n=107) compared with healthy controls (HCs) (n=140), adjusting for age, sex, scan site and whole AH volumes.

|  | **Mixed effect size estimates d**  **(SAD vs HCs)** | **Standard error of *d*** | **FDR corrected**  **p value** | **Uncorrected p-value** |  | **Mixed effect size estimates d**  **(SAD vs HCs)** | **Standard error of *d*** | **FDR corrected**  **p value** | **Uncorrected p-value** |
| --- | --- | --- | --- | --- | --- | --- | --- | --- | --- |
| L Lateral | -0.16 | 0.13 | 0.39 | 0.18 | R Lateral | -0.01 | 0.13 | 0.94 | 0.94 |
| L Basal | -0.20 | 0.13 | 0.26 | 0.09 | R Basal | -0.11 | 0.13 | 0.54 | 0.36 |
| L Accessory Basal | -0.29 | 0.13 | 0.08 | 0.01 | R Accessory Basal | -0.21 | 0.13 | 0.26 | 0.09 |
| L AAA | 0.01 | 0.13 | 0.94 | 0.94 | R AAA | -0.06 | 0.13 | 0.76 | 0.62 |
| L Central | -0.09 | 0.13 | 0.60 | 0.44 | R Central | -0.24 | 0.13 | 0.17 | 0.04 |
| L Medial | -0.21 | 0.13 | 0.20 | 0.06 | R Medial | -0.10 | 0.13 | 0.54 | 0.38 |
| L Cortical | -0.24 | 0.13 | 0.15 | 0.03 | R Cortical | -0.03 | 0.13 | 0.89 | 0.80 |
| L Corticoamygdaloid transition | -0.36 | 0.13 | 0.06 | 0.00 | R Corticoamygdaloid transition | -0.10 | 0.13 | 0.60 | 0.44 |
| L Paralaminar | -0.18 | 0.13 | 0.37 | 0.17 | R Paralaminar nucleus | 0.11 | 0.13 | 0.54 | 0.37 |
| L Parasubiculum | 0.03 | 0.13 | 0.89 | 0.83 | R Parasubiculum | 0.12 | 0.13 | 0.54 | 0.34 |
| L Subiculum | 0.13 | 0.13 | 0.51 | 0.30 | R Subiculum | 0.25 | 0.13 | 0.17 | 0.05 |
| L presubiculum | -0.05 | 0.13 | 0.76 | 0.64 | R Presubiculum | 0.06 | 0.13 | 0.76 | 0.65 |
| L CA1 | 0.06 | 0.13 | 0.76 | 0.61 | R CA1 | 0.15 | 0.13 | 0.44 | 0.23 |
| L CA3 | 0.35 | 0.13 | 0.06 | 0.01 | R CA3 | 0.34 | 0.13 | 0.06 | 0.01 |
| L CA4 | 0.36 | 0.13 | 0.06 | 0.01 | R CA4 | 0.40 | 0.13 | 0.06 | 0.00 |
| L GC ML DG | 0.30 | 0.13 | 0.10 | 0.02 | R GC ML DG | 0.33 | 0.13 | 0.06 | 0.01 |
| L Molecular layer | 0.15 | 0.13 | 0.44 | 0.23 | R Molecular layer | 0.19 | 0.13 | 0.30 | 0.13 |
| L Fimbria | 0.02 | 0.13 | 0.89 | 0.85 | R Fimbria | -0.03 | 0.13 | 0.89 | 0.82 |
| L HATA | -0.23 | 0.13 | 0.17 | 0.05 | R HATA | -0.13 | 0.13 | 0.48 | 0.27 |
| L Hippocampal fissure | 0.06 | 0.13 | 0.76 | 0.60 | R Hippocampal fissure | 0.19 | 0.13 | 0.30 | 0.13 |
| L Hippocampal tail | 0.17 | 0.13 | 0.30 | 0.13 | R Hippocampal tail | 0.13 | 0.13 | 0.45 | 0.25 |

Abbreviations: cornu ammonis (CA) sectors, CA1, CA2-3, CA4, granule cell layer of dentate gyrus (DG), molecular layer (ML), hippocampus–amygdala transition area (HATA), corticoamygdaloid transition area (CAT), anterior amygdaloid area (AAA)

**Supplementary table S2:** Mega analysis linear regression results for **bilateral** amygdala and hippocampal subfield volumes for **patients with SAD without comorbid anxiety disorder** (n=86) compared with adult healthy controls (HCs) (n=140), adjusting for age, sex, scan site and whole AH volumes.

|  | **Mixed effect size estimates d**  **(SAD vs HCs)** | **Standard error of *d*** | **FDR corrected**  **p value** | **Uncorrected p.value** |
| --- | --- | --- | --- | --- |
| Lateral | -0.305 | 0.134 | 0.037 | 0.016 |
| Basal | -0.335 | 0.135 | 0.020 | 0.006 |
| Accessory Basal | -0.451 | 0.135 | 0.005 | 0.000 |
| AAA | -0.111 | 0.134 | 0.455 | 0.368 |
| Central | -0.246 | 0.134 | 0.073 | 0.035 |
| Medial | -0.162 | 0.134 | 0.217 | 0.145 |
| Cortical | -0.191 | 0.134 | 0.144 | 0.083 |
| Corticoamygdaloid transition | -0.424 | 0.135 | 0.007 | 0.001 |
| Paralaminar | -0.125 | 0.134 | 0.455 | 0.355 |
| Parasubiculum | 0.070 | 0.134 | 0.665 | 0.601 |
| Presibiculum | -0.040 | 0.134 | 0.786 | 0.748 |
| Subiculum | 0.196 | 0.134 | 0.212 | 0.131 |
| CA1 | 0.025 | 0.134 | 0.839 | 0.839 |
| CA3 | 0.376 | 0.135 | 0.020 | 0.006 |
| CA4 | 0.472 | 0.135 | 0.007 | 0.001 |
| GC ML DG | 0.355 | 0.135 | 0.023 | 0.008 |
| Molecular layer | 0.336 | 0.135 | 0.018 | 0.004 |
| HATA | -0.330 | 0.135 | 0.027 | 0.010 |
| Fimbria | -0.106 | 0.134 | 0.490 | 0.420 |
| Hippocampal fissure | 0.144 | 0.134 | 0.384 | 0.274 |
| Hippocampal tail | 0.207 | 0.134 | 0.140 | 0.073 |

Abbreviations: cornu ammonis (CA) sectors, CA1, CA2-3, CA4, granule cell layer of dentate gyrus (DG), molecular layer (ML), hippocampus–amygdala transition area (HATA), corticoamygdaloid transition area (CAT), anterior amygdaloid area (AAA).

**Supplementary table S3:** Mega analysis linear regression results for **bilateral** amygdala and hippocampal subfield volumes for **patients with SAD without MDD** (n=83) compared with adult healthy controls (HCs) (n=140), adjusting for age, sex, scan site and whole AH volumes.

|  | **Mixed effect size estimates d**  **(SAD vs HCs)** | **Standard error of *d*** | **FDR corrected**  **p value** | **Uncorrected p.value** |
| --- | --- | --- | --- | --- |
| Lateral | -0.196 | 0.135 | 0.194 | 0.119 |
| Basal | -0.294 | 0.135 | 0.085 | 0.016 |
| Accessory Basal | -0.407 | 0.136 | 0.017 | 0.001 |
| AAA | -0.148 | 0.135 | 0.358 | 0.239 |
| Central | -0.253 | 0.135 | 0.102 | 0.030 |
| Medial | -0.173 | 0.135 | 0.194 | 0.120 |
| Cortical | -0.227 | 0.135 | 0.104 | 0.040 |
| Corticoamygdaloid transition | -0.326 | 0.135 | 0.085 | 0.012 |
| Paralaminar | -0.130 | 0.135 | 0.483 | 0.345 |
| Parasubiculum | 0.051 | 0.135 | 0.788 | 0.702 |
| Presibiculum | -0.022 | 0.135 | 0.889 | 0.863 |
| Subiculum | 0.091 | 0.135 | 0.634 | 0.483 |
| CA1 | 0.046 | 0.135 | 0.788 | 0.713 |
| CA3 | 0.290 | 0.135 | 0.102 | 0.034 |
| CA4 | 0.363 | 0.136 | 0.085 | 0.008 |
| GC ML DG | 0.244 | 0.135 | 0.128 | 0.061 |
| Molecular layer | 0.257 | 0.135 | 0.097 | 0.023 |
| HATA | -0.255 | 0.135 | 0.111 | 0.048 |
| Fimbria | -0.019 | 0.135 | 0.889 | 0.889 |
| Hippocampal fissure | 0.081 | 0.135 | 0.664 | 0.538 |
| Hippocampal tail | 0.192 | 0.135 | 0.189 | 0.099 |

Abbreviations: cornu ammonis (CA) sectors, CA1, CA2-3, CA4, granule cell layer of dentate gyrus (DG), molecular layer (ML), hippocampus–amygdala transition area (HATA), corticoamygdaloid transition area (CAT), anterior amygdaloid area (AAA).

**Supplementary table S4:** Mega analysis linear regression results for **bilateral** amygdala and hippocampal subfield volumes for **patients with SAD without medication** (n=89) compared with adult healthy controls (HCs) (n=140), adjusting for age, sex, scan site and whole AH volumes.

|  | **Mixed effect size estimates d**  **(SAD vs HCs)** | **Standard error of *d*** | **FDR corrected**  **p value** | **Uncorrected p.value** |
| --- | --- | --- | --- | --- |
| Lateral | -0.251 | 0.133 | 0.127 | 0.048 |
| Basal | -0.288 | 0.133 | 0.083 | 0.020 |
| Accessory Basal | -0.362 | 0.134 | 0.063 | 0.003 |
| AAA | -0.101 | 0.133 | 0.502 | 0.407 |
| Central | -0.202 | 0.133 | 0.169 | 0.081 |
| Medial | -0.120 | 0.133 | 0.395 | 0.282 |
| Cortical | -0.129 | 0.133 | 0.347 | 0.230 |
| Corticoamygdaloid transition | -0.312 | 0.134 | 0.080 | 0.015 |
| Paralaminar | -0.134 | 0.133 | 0.423 | 0.322 |
| Parasubiculum | 0.061 | 0.133 | 0.705 | 0.638 |
| Presibiculum | -0.026 | 0.133 | 0.830 | 0.830 |
| Subiculum | 0.197 | 0.133 | 0.227 | 0.130 |
| CA1 | 0.028 | 0.133 | 0.830 | 0.823 |
| CA3 | 0.276 | 0.133 | 0.120 | 0.040 |
| CA4 | 0.358 | 0.134 | 0.067 | 0.009 |
| GC ML DG | 0.245 | 0.133 | 0.144 | 0.062 |
| Molecular layer | 0.257 | 0.133 | 0.096 | 0.027 |
| HATA | -0.327 | 0.134 | 0.067 | 0.010 |
| Fimbria | -0.084 | 0.133 | 0.603 | 0.517 |
| Hippocampal fissure | 0.156 | 0.133 | 0.347 | 0.231 |
| Hippocampal tail | 0.177 | 0.133 | 0.227 | 0.120 |

Abbreviations: cornu ammonis (CA) sectors, CA1, CA2-3, CA4, granule cell layer of dentate gyrus (DG), molecular layer (ML), hippocampus–amygdala transition area (HATA), corticoamygdaloid transition area (CAT), anterior amygdaloid area (AAA).

**Supplementary table S5:** Mega analysis linear regression results for **bilateral** amygdala and hippocampal subfield volumes and **symptom severity** (n=76) adjusting for age, sex, and scan site whole AH volumes.

|  | **rME (SAD vs HCs)** | **Standard error of *rME*** | **FDR corrected**  **p value** | **Uncorrected p.value** |
| --- | --- | --- | --- | --- |
| Lateral | -0.138 | 0.117 | 0.809 | 0.249 |
| Basal | 0.000 | 0.117 | 0.998 | 0.998 |
| Accessory Basal | 0.051 | 0.117 | 0.809 | 0.616 |
| AAA | -0.076 | 0.117 | 0.809 | 0.448 |
| Central | 0.089 | 0.117 | 0.809 | 0.458 |
| Medial | 0.103 | 0.117 | 0.809 | 0.313 |
| Cortical | 0.054 | 0.117 | 0.809 | 0.613 |
| Corticoamygdaloid transition | -0.015 | 0.117 | 0.998 | 0.879 |
| Paralaminar | -0.125 | 0.117 | 0.809 | 0.297 |
| Parasubiculum | -0.064 | 0.117 | 0.809 | 0.578 |
| Presibiculum | -0.091 | 0.117 | 0.809 | 0.439 |
| Subiculum | -0.016 | 0.117 | 0.998 | 0.887 |
| CA1 | 0.061 | 0.117 | 0.809 | 0.602 |
| CA3 | 0.108 | 0.117 | 0.809 | 0.354 |
| CA4 | 0.147 | 0.117 | 0.809 | 0.208 |
| GC ML DG | 0.114 | 0.117 | 0.809 | 0.342 |
| Molecular layer | 0.097 | 0.117 | 0.809 | 0.392 |
| HATA | -0.062 | 0.117 | 0.809 | 0.543 |
| Fimbria | 0.007 | 0.117 | 0.998 | 0.950 |
| Hippocampal fissure | 0.088 | 0.117 | 0.809 | 0.464 |
| Hippocampal tail | -0.004 | 0.117 | 0.998 | 0.961 |

Abbreviations: cornu ammonis (CA) sectors, CA1, CA2-3, CA4, granule cell layer of dentate gyrus (DG), molecular layer (ML), hippocampus–amygdala transition area (HATA), corticoamygdaloid transition area (CAT), anterior amygdaloid area (AAA).

|  | **SAD with comorbid (lifetime) anxiety (n=31)** | **SAD with comorbid (lifetime) MDD (n=24)** |
| --- | --- | --- |
| Lifetime comorbid ANX (n, %) | 10 (32.25%) | - |
| Current comorbid ANX (n, %) | 21 (67.74%) | - |
| Lifetime comorbid MDD (n, %) | - | 19 (79.16%) |
| Current comorbid MDD (n, %) | - | 5 (20.83%) |
| Medication use at scan (n, %) | 4 (22.22%) | 6 (33.33%) |

**Supplementary table S6:** Clinical characteristics for samples included in the sub-group mega-analysis (reported for individuals with SAD).

Note: Medication use at scan (n=18).

**Supplementary table S7:** SAD and HCs left and right mean values for amygdala and hippocampal subfields

Mega analysis linear regression results for bilateral amygdala and hippocampal subfield volumes for patients with SAD (n=107) compared with healthy controls (HCs) (n=140), adjusting for age, sex, scan site, left whole HA and right whole HA.

| **L-Amygdala subfields** | **L-mean value** | **L-pFDR** | **R-mean value** | **R-pFDR** |
| --- | --- | --- | --- | --- |
| Lateral | 664.48 | 0.39 | 695.83 | 0.94 |
| Basal | 470.84 | 0.26 | 482.88 | 0.54 |
| Accessory Basal | 293.27 | 0.08 | 297.94 | 0.26 |
| AAA | 59.22 | 0.94 | 64.94 | 0.76 |
| Central | 47.68 | 0.60 | 54.13 | 0.17 |
| Medial | 24.27 | 0.20 | 25.63 | 0.54 |
| Cortical | 29.58 | 0.15 | 29.22 | 0.89 |
| Corticoamygdaloid transition | 203.41 | 0.06 | 198.51 | 0.60 |
| Paralaminar | 52.50 | 0.37 | 51.71 | 0.54 |

| **L-Hippocampal subfields** | **L-mean value** | **L-pFDR** | **R-mean value** | **R-pFDR** |
| --- | --- | --- | --- | --- |
| Parasubiculum | 66.42 | 0.89 | 66.56 | 0.54 |
| Presubiculum | 301.45 | 0.51 | 282.66 | 0.76 |
| Subiculum | 436.92 | 0.76 | 428.41 | 0.17 |
| CA1 | 641.57 | 0.76 | 677.95 | 0.44 |
| CA3 | 216.75 | 0.06 | 235.58 | 0.06 |
| CA4 | 250.24 | 0.06 | 261.55 | 0.06 |
| GC ML DG | 291.34 | 0.10 | 302.71 | 0.06 |
| Molecular layer HP | 557.70 | 0.44 | 575.43 | 0.30 |
| HATA | 66.02 | 0.89 | 66.81 | 0.48 |
| Fimbria | 97.47 | 0.17 | 92.49 | 0.30 |
| Hippocampal fissure | 147.06 | 0.76 | 151.28 | 0.89 |
| Hippocampal tail | 527.11 | 0.30 | 525.50 | 0.45 |

**Supplementary table S8:** SAD left and right mean values for amygdala and hippocampal subfields

| **L-Amygdala subfields** | **L-mean value** | **R-mean value** |
| --- | --- | --- |
| Lateral | 646.61 | 682.52 |
| Basal | 456.29 | 470.44 |
| Accessory Basal | 278.93 | 286.85 |
| AAA | 58.02 | 63.16 |
| Central | 44.97 | 49.96 |
| Medial | 21.82 | 23.58 |
| Cortical | 27.32 | 27.95 |
| Corticoamygdaloid transition | 193.98 | 194.19 |
| Paralaminar | 51.51 | 51.55 |

| **L-Hippocampal subfields** | **L-mean value** | **R-mean value** |
| --- | --- | --- |
| Parasubiculum | 66.90 | 67.18 |
| Presubiculum | 302.22 | 284.13 |
| Subiculum | 438.69 | 431.09 |
| CA1 | 641.29 | 680.33 |
| CA3 | 220.62 | 239.14 |
| CA4 | 252.50 | 264.70 |
| GC ML DG | 293.51 | 305.38 |
| Molecular layer HP | 560.48 | 578.16 |
| HATA | 64.40 | 65.84 |
| Fimbria | 96.86 | 91.46 |
| Hippocampal fissure | 145.19 | 152.41 |
| Hippocampal tail | 533.06 | 526.35 |

| **L-Amygdala subfields** | **L-mean value** | **R-mean value** |
| --- | --- | --- |
| Lateral | 664.48 | 695.83 |
| Basal | 470.84 | 482.88 |
| Accessory Basal | 293.27 | 297.94 |
| AAA | 59.22 | 64.94 |
| Central | 47.68 | 54.13 |
| Medial | 24.27 | 25.63 |
| Cortical | 29.58 | 29.22 |
| Corticoamygdaloid transition | 203.41 | 198.51 |
| Paralaminar | 52.50 | 51.71 |

**Supplementary table S9:** HCs left and right mean values for amygdala and hippocampal subfields

| **L-Hippocampal subfields** | **L-mean value** | **R-mean value** |
| --- | --- | --- |
| Parasubiculum | 66.05 | 66.09 |
| Presubiculum | 300.86 | 281.54 |
| Subiculum | 435.58 | 426.36 |
| CA1 | 641.79 | 676.14 |
| CA3 | 213.80 | 232.85 |
| CA4 | 248.52 | 259.14 |
| GC ML DG | 289.69 | 300.67 |
| Molecular layer HP | 555.57 | 573.34 |
| HATA | 67.25 | 67.55 |
| Fimbria | 97.93 | 93.28 |
| Hippocampal fissure | 148.50 | 150.42 |
| Hippocampal tail | 522.57 | 524.86 |
